# Inhibition-Dominated Rich-Club Shapes Dynamics in Cortical Microcircuits

**DOI:** 10.1101/2021.05.07.443074

**Authors:** Hadi Hafizi, Sunny Nigam, Josh Barnathan, Naixin Ren, Ian H Stevenson, Sotiris C Masmanidis, Ehren L Newman, Olaf Sporns, John M Beggs

**Affiliations:** Department of Psychological and Brain Sciences, Indiana University, Bloomington, Indiana, 47405, USA; Department of Neurobiology and Anatomy, McGovern Medical School, University of Texas, Houston, Texas, 77030, USA; Department of Physics, Indiana University, Bloomington, Indiana, USA 47405; Department of Psychological Sciences, University of Connecticut, Storrs, Connecticut, 06269, USA; Department of Biomedical Engineering, University of Connecticut, Storrs, Connecticut, 06030, USA; Department of Neurobiology, David Geffen School of Medicine, University of California, Los Angeles, California, 90095, USA; Indiana University Network Science Institute, Indiana University, Bloomington, Indiana, 47408, USA

## Abstract

Functional networks of cortical neurons contain highly interconnected hubs, forming a rich-club structure. However, the cell type composition within this distinct subnetwork and how it influences large-scale network dynamics is unclear. Using spontaneous activity recorded from hundreds of cortical neurons in orbitofrontal cortex of awake behaving mice and from organotypic cultures, we show that the rich-club is disproportionately composed of inhibitory neurons, and that inhibitory neurons within the rich-club are significantly more synchronous than other neurons. At the population level, neurons in the rich-club exert higher than expected Granger causal influence on overall population activity at a broad range of frequencies compared to other neurons. Finally, neuronal avalanche duration is significantly correlated with the fraction of rich neurons that participate in the avalanche. Together, these results suggest an unexpected role of a highly connected, inhibition-rich subnetwork in driving and sustaining activity in local cortical networks.

**SIGNIFICANCE STATEMENT:** It is widely believed that the relative abundance of excitatory and inhibitory neurons in cortical circuits is roughly 4:1. This relative abundance has been widely used to construct numerous cortical network models. Here we show that contrary to this notion, a sub-network of highly connected hub neurons (rich-club) consists of a higher abundance of inhibitory neurons compared to that found in the entire network or the non-rich subnetwork. Inhibitory hub neurons contribute to higher synchrony within the rich club compared to the rest of the network. Strikingly, higher activation of the inhibition-dominated rich club strongly correlates with longer avalanches in cortical circuits. Our findings reveal how network topology combined with cell-type specificity orchestrates population wide activity in cortical microcircuits.

## INTRODUCTION

Connectivity in brains is far from egalitarian. Some regions in the human cortex send and receive fiber bundles at a density ten times greater than others (Hagmann et al., 2008; Markov et al., 2013); the top 20% of cortical neurons in mouse carry 70% of the information flow (Nigam et al., 2016); the nervous system of the worm *C. elegans* contains a small number of well-connected neurons (Towlson et al., 2013). Even less democratic is the finding that in each of these systems, the most highly connected units (cortical regions, hub neurons) connect to each other more than expected by chance, forming what is called a rich-club structure (Dann et al., 2016; van den Heuvel and Sporns, 2011; Nigam et al., 2016; Towlson et al., 2013). Despite the ubiquity of the rich-club network structure across different species, the cell type composition within this distinct subnetwork and how it shapes network dynamics is unknown.

In the past decade, the relationship between neuron types and network structure has received increased research attention. In the developing hippocampus, hub neurons are all inhibitory (Bonifazi et al., 2009); more recent work in mouse somato-motor cortex slices found inhibitory neurons were more topologically central within networks than excitatory neurons (Kajiwara et al., 2021). However, neither of these studies directly examined the rich club and its cell type composition. Hence, it is unknown whether the rich-club is composed of exclusively excitatory neurons, or if its composition corresponds to typically reported values (85%) of excitatory neurons in cortical networks (Douglas and Martin, 2004).

More importantly, the influence of the rich-club on dynamics in local cortical networks is poorly understood. Recordings from multiple motor areas in behaving monkeys showed that the most highly connected rich-club neurons, which spanned these areas, were synchronous in beta and low frequency bands (Dann et al., 2016). Synchrony is thought to be a mechanism for forming assemblies of neurons and for coordinating switching between them (Cho et al., 2020; Fries, 2005; Tort et al., 2007). Recent modeling studies suggest that the rich-club could initiate and sustain cortical activity (Aguilar-Velázquez and Guzmán-Vargas, 2019; Gu et al., 2019), but this has not been experimentally observed yet.

To investigate these issues, we analyzed dense electrode array recordings collected from awake behaving mice and from cortical slice cultures; the rich club had been previously reported in both data sets (Nigam et al., 2016). Despite the substantial differences in these preparations, we found that they were qualitatively similar in several respects. First, for both, the proportion of inhibitory neurons within the rich-club was significantly higher than outside the rich club or the entire network. Second, inhibitory neurons within the rich-club were more synchronous compared to other cell types found within and outside the rich-club for both preparations. Third, rich club neurons exerted higher Granger causal influence on the rest of the network compared to non-rich neurons. Finally, the proportion of rich club neurons active during neuronal avalanches showed a significant positive correlation with avalanche length. Together, these findings suggest that the rich club is rich in inhibition and plays a central role in shaping dynamics in local cortical networks. Portions of this work were previously presented in thesis form (Hafizi, 2020).

## METHODS

### Experimental design

All in-vivo recording procedures were approved by the University of California, Los Angeles, Chancellor’s Animal Research Committee. Data was recorded from single housed male mice C57/B1/6J (N = 7, 12-16 weeks old; The Jackson Laboratory) using silicon microprobes as described in a previous study (Shobe et al., 2015). Briefly, extra-cellular activity of hundreds of neurons was recorded simultaneously (sampled at 25 kHz) from awake behaving mice using a 5 shank, 256 site silicon microprobe (50-54 sites per shank arranged in a hexagonal pattern, ∼30 μm inter-site spacing, and 300-400 μm inter-shank spacing) inserted into the orbitofrontal cortex. Consecutive microprobes were separated by roughly 0.3 -0.4 mm. Spike sorting was performed offline using a semi-automated Matlab script. Briefly, the 1024 electrodes were subdivided into groups of 2-5 local electrodes. Due to the high density of electrodes, a single neuron was detected on multiple channels. The mean background signal from all electrodes on a silicon prong was subtracted from the signal of each electrode to minimize globally correlated signal artifacts. Signals were then filtered between 600-6500 Hz, and spikes were identified on the spike trough with an SNR of ∼ 3. Putative units were then isolated on these local electrodes. Units were isolated by automatically generating waveform-based templates (Rutishauser & Mamelak 2006) using data from the first 5 mins of the recording session and then matching the waveforms recorded in the rest of the session. Duplicate units on nearby electrodes were identified by comparing mean spike waveform and discarded. Time stamps from the remaining units were used to repeat waveform collection under a wider band (300 – 6500 Hz) for visualizing and manually scoring the units. Units were manually inspected and removed if there were signal artifacts, minimum waveform amplitude was lower than 50 μV or if there were fewer than 150 spikes during the entire recording session. The data used for this analysis were prepared by concatenating resting period activity in between periods when the mouse was performing an odor discrimination task in the same recording session. Resting corresponded to periods of immobility (no treadmill movement and no licking) and a lack of explicitly presented task related stimuli. The firing rates were fairly constant across resting periods except for the last 2 s, where a consistent increase in firing rate was detected. To avoid possible non-stationarities, we deleted the last 2 s from each resting period and then concatenated the different resting periods identified across the entire recording session together to obtain the final spike trains for each recorded neuron.

Cultured tissue from animals was prepared according to guidelines from the National Institutes of Health, and all animal procedures were approved by the Animal Care and Use Committees at Indiana University and at the University of California, Santa Cruz. An extended description of the *in-vitro* data collection can be found in (Friedman et al. 2012). Briefly, brains from 6 mouse pups (postnatal day 6 to 7; RRID:Charles_River:24101632, Harlan) were removed under a sterile hood and placed in chilled Gey’s balanced salt solution (Sigma-Aldrich) for 1 h at 0°C. After 30 min, half the solution was changed and brains were next blocked into 5 mm^2^ sections containing the somatosensory cortex (Paxinos and Watson 1986). Blocks were then sliced with a thickness of 400 μm using a vibrating blade microtome (Leica VT1000 S). Cultures that are prepared using this method are called “organotypic” slice cultures. They have been shown to preserve many of the organ’s features and they produce a variety of emergent network activity patterns that have been found *in-vivo*, including oscillations (Baker, Corner, and van Pelt 2006; Gireesh and Plenz 2008), synchrony (Beggs and Plenz 2004; Baker, Corner, and van Pelt 2006), and neuronal avalanches (Beggs and Plenz 2003; Friedman et al. 2012). After 2-3 weeks in the incubator, the somatosensory cortex portion of slice cultures were placed on top of a 512 micro-electrode array (MEA) while their spontaneous electrical activity was recorded. The electrodes are arranged as a triangular lattice in a rectangular space of 1.8 mm x 0.9 mm. This recording system has been previously used to record spikes in the retina (Litke et al. 2004; Shlens et al. 2006; Petrusca et al. 2007; Field et al. 2010) and in the cortex (Tang et al. 2008; Friedman et al. 2012). The 512-MEA records electrical activity at 20 kHz and the electrode spacing is 60 μm. This means the most temporally detailed neuronal activity could be recorded from up to about 500 neurons simultaneously. Recording sessions lasted for 1.5 hours and we discarded the first half hour to exclude possible transients. Finally, spike sorting was performed to identify spikes belonging to individual neurons. The spike sorting process has previously been described in (Litke et al. 2004; Tang et al. 2008).

### Cell-type classification/identification

Classification of cell types was performed by following methods reported in a recent study (Ren et al., 2020). Briefly, a Gaussian mixture model was fitted to features of the extracellularly recorded spike waveforms (Bishop, 2006). This unsupervised clustering method is computationally inexpensive and provides a relatively accurate indicator of cell-type without a priori knowledge of the ground truth (e.g., from histological analysis on the tissue or optotagging) (Kim et al., 2016; Lima et al., 2009). We calculated three parameters of the spike waveform used in previous studies in identifying cell types (Barthó et al., 2004; Sirota et al., 2008; Stark et al., 2013; Trainito et al., 2019): trough-to-peak duration, half amplitude duration and logarithm of firing rate. Trough-to-peak duration is defined as the time interval between the global minimum of the spike waveform and the following local maximum. Half amplitude duration is the duration between the two points where the spike waveform crosses amplitude half the peak. Finally, firing rate is the average number of spikes per second. Although correlated, the first two measures capture different aspects of the intracellular action potential: the speed of depolarization and of the subsequent after-hyperpolarization (Henze et al., 2000) and are both distinguishing features of neuronal cell types (Nowak et al., 2003). Using these three features, we fit the parameters of a 2-component mixture model (means, covariance matrices, and prior probabilities) using an Expectation Maximization algorithm. We then calculated the posterior probability of each neuron being excitatory or inhibitory. Here we chose a conservative threshold of 90% likelihood to label neurons as excitatory or inhibitory.

### Transfer Entropy analysis

Transfer Entropy (TE) (Schreiber, 2000) was used to quantify effective connectivity between neurons using the Transfer Entropy Toolbox (Ito et al., 2011) as was described in our previous study (Nigam et al., 2016). TE is an asymmetric information theoretic measure that quantifies causal, non-linear interactions between source and target neurons. TE is non-zero if inclusion of the past activity of the source neuron improves the prediction of the spiking of the target neuron beyond the prediction from the past spiking of the target neuron itself. Briefly, we calculated time lagged TE at a range of delays (1-30ms) between the spiking activity of simultaneously recorded pairs of neurons (source and target). To control for firing rate, we calculated TE between the pairs when the spike times of the target neuron were jittered by 1-19 ms drawn from a normal distribution. The mean of the jittered value was then subtracted from the raw estimate between the pairs. Thus, the TE values that remained were directly caused by timing relationships at a resolution less than 20 ms. To control for overall network drive which could lead to increased occurrences of spike coincidences between pairs of neurons, we only considered pairs where TE as a function of time lag had a sharp peak. This was quantified by calculating Coincidence Index (CI) (Ito et al., 2011; Nigam et al., 2016; Shimono and Beggs, 2015) where we evaluated the ratio of the area around the peak (peak time ± 4 ms) to that of the area under the entire TE vs time lag profile (0-30 ms). Only pairs with both high CI (sharply peaked) and TE were considered for the analysis (see also Shimono and Beggs, 2015). The threshold for peak TE and CI was chosen based on TE analysis performed on spiking data generated from a cortical network model where the synaptic connectivity was known beforehand. Values of CI and TE_peak_ that maximized the ratio of true positive rate (TPR) to false positive rate (FPR) for the network model were used to threshold the effective connectivity estimated from the in-vivo recordings. Finally, possible spurious connections arising out of common drive and transitive drive were eliminated based on analyzing delays at which significant TE was detected in neuronal pairs. This effective connectivity analysis generated sparse, weighted and directed graphs between hundreds of simultaneously recorded neurons.

### Rich-club analysis

To quantify the strength of connections within subnetworks, we used the weighted normalized rich-club coefficient (Colizza et al., 2006; Opsahl et al., 2008) as described in detail in (Nigam et al., 2016). Briefly, we defined the richness parameter *r* of each neuron as the sum of the total outgoing and incoming TE from and into that neuron, respectively (Figure 2A). Then a list of the unique values of the richness parameter was created, ranked from smallest to largest (*r*_*min*_, *r*_*2*_, … *r*_*max*_). Additionally, a list of the pairwise TE values (TE_rank_) ranked from largest to smallest was also created (TE_max_, TE_2_, … TE_min_). Next, we isolated the subnetwork where all neurons had a richness parameter > *r*_*k*_ and counted the number of edges between neurons and defined it as 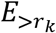. We then summed all the pairwise TE values in that subnetwork and defined it to be 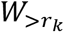. The weighted rich-club coefficient 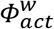 is the ratio of 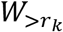 to the sum of the 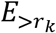 strongest pairwise TE values in the network obtained from the list (TE_rank_). This analysis generated weighted rich-club coefficients for each value of the ranked richness parameters (*r*_*min*_, *r*_*2*_, … *r*_*max*_) defined above. This ratio represents what fraction of the strongest weights in the whole network is present in the subnetwork. To examine whether these coefficients were any different from what would be expected by chance, we used the Brain Connectivity toolbox to generate 1000 randomized versions of the actual networks such that the richness parameters of the neurons were unchanged in each randomized network (Rubinov and Sporns, 2010). We then calculated the weighted rich-club coefficients at the same thresholds from the randomized networks 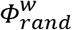. The normalized richness coefficient 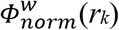 at each richness parameter was defined as the ratio of 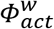 to 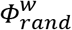. If 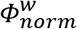 was significantly greater than 1 (Wilcoxon signed rank test; P < 0.05) for a range of the richness parameter values, then a rich-club existed in that regime. To correct for multiple comparisons over the range of the richness parameters, false discovery rate (FDR) correction (Benjamini and Yekutieli, 2001) was implemented, limiting the FDR to 0.05.

### Cross-correlation analysis

We used the Transfer Entropy Toolbox (Ito et al., 2011), to calculate Normalized Cross-Correlation Histograms (NCCH) for pairs of binary spike trains (Brosch and Schreiner, 1999) as follows:

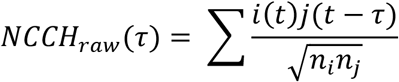

where *i*(*t*), *j*(*t*) are the binary states of the neurons at time *t*, i.e., either 1 or 0 based on whether the neurons fired an action potential. *τ* is the positive or negative lag in milliseconds at which the state of the second neuron in the pair is evaluated. *n*_*i*_ and *n*_*j*_ are the total number of spikes fired by neuron *i* and *j* respectively. The denominator represents the normalization factor which is the geometric mean of the number of spikes fired by each neuron constituting the pair. Shuffled estimates of NCCH were generated by spike jittering where each spike time was shifted by +/-*t*_*shuf*_ drawn randomly from a uniform distribution with 0 mean and a width of 30 ms. The shuffling procedure jittered the temporal pattern of activity of the two neurons but preserved the total number of spikes fired by each neuron. The NCCH values used throughout the analysis were obtained by subtracting the shuffled estimates from the raw values. This correction accounts for correlations arising between spike trains just due to higher firing rates instead of a specific temporal structure at short time scales.

### Synchrony Index (SI)

We defined the Synchrony Index as the area under the shuffle corrected NCCH curve centered at 0 lag and extending to 30 ms on either side (see Figure 3A). Higher values of SI indicate more coincident activity between neuronal pairs within a short time interval and hence more pairwise synchrony. We calculated SI for all pairs of neurons in the data sets and partitioned them into 3 classes: rich-rich, rich-nonrich and nonrich-nonrich based on the membership of the neurons to the rich and non-rich subnetwork. Based on the cell type classification described above we further divided pairs within the rich and non-rich subnetworks into excitatory-excitatory, inhibitory-inhibitory and excitatory-inhibitory pairs and evaluated SI for each of these categories of pairs.

### Population Normalized Population Cross-Correlation (NPCCH)

In order to quantify how the activity of individual neurons influenced the dynamics of specific subnetworks, we calculated the shuffle corrected normalized correlation histogram between the spike train of a reference neuron and all the spikes fired by the rest of the neurons in a defined sub-population (not including the reference neuron) (Okun et al., 2015). The normalization constant in this case was the geometric mean of the total number of spikes of the reference neuron and that of the sub-population of neurons. Similar to the analysis for pairs, the population measure was evaluated at different positive and negative time lags (-150 ms to +150 ms) with respect to each spike of the reference neuron. Raw values of the NPCCH were corrected by subtracting the shuffled estimates obtained by shuffling the spike times of the reference neuron and that of neurons in the subnetwork with the same jitter paradigm used for the pairwise analysis. Area under the NPCCH was evaluated for a pre-period extending from -30 to 0 ms and a post period extending from 0 to +30 ms. 0 ms represents the time point at which the reference neuron fires.

### Granger Causality

Granger Causality (GC) was used to quantify the influence of population spiking activity in one sub-network on the other. Specifically, we generated two new “population” time-series by summing all the spikes from neurons belonging to each subnetwork respectively. The Akaike information criterion (Akaike, 1974), a principled way to determine the number of time steps included, was then used to calculate the order up to 250 time-bins, and GC was calculated using the ‘Multivariate Granger Causality toolbox’ (MVGC, Barnett and Seth, 2014). To calculate GC between two random subnetworks, we randomly sampled two populations that were size-matched to the rich and non-rich subnetwork respectively 50 times, and calculated shuffle corrected GC between these subnetworks. We then subtracted the GC estimates for the random subnetworks from the estimates calculated for the actual rich and non-rich subnetworks ΔGC (actual – random; Figure 4 K, L). A signed rank test followed by Bonferroni-Holm test for multiple comparisons was used to test for significance of the deviation of ΔGC values from 0 for each frequency value.

### Avalanche dynamics

In order to identify the role of connectivity and cell types in sustaining and controlling activity in neural circuits, we examined their participation across time in neuronal avalanches. Avalanches are defined as consecutive time-bins of neural activity with at least one spike bracketed by time-bins with no spikes (Figure 5A) and have been reported in both *in-vitro* (Beggs and Plenz, 2003, 2004; Rolston et al., 2007) and *in-vivo* (Hahn et al., 2010; Petermann et al., 2009). Avalanche length (L) was defined as the number of consecutive time bins with at least one active neuron. To further investigate the role of neurons and their cell type in avalanches, we examined the proportion of active neurons within and outside of the rich-club with respect to all active neurons in every time bin along avalanches of different lengths.

## RESULTS

We analyzed spiking activity in the orbitofrontal cortex of awake mice (n = 990 neurons; N = 7 mice) with silicon microprobes (Shobe et al., 2015) (Figure 1A). In addition, we also analyzed spiking activity recorded in organotypic cultures (coronal cortical slices, n = 4768 neurons; N = 14 sessions; Figure 1B). For recordings in awake mice, we limited our analysis to portions of the data when the mice were not involved in a task and were immobile on the treadmill (see Materials and Methods). In a prior study (Nigam et al., 2016), we used Transfer Entropy (TE) to construct effective connectivity networks (weighted and directed graphs) between hundreds of neurons in each session (both in-vivo and in-vitro). We discovered that these networks contained a distinct sub-network of hub neurons (high incoming and outgoing TE connections) with a higher-than-expected density and strength of connections among them forming a rich-club (Figure 1C) (Nigam et al., 2016). Neurons outside the rich-club formed the non-rich sub-network characterized by fewer and weaker connections.

**Figure 1.**
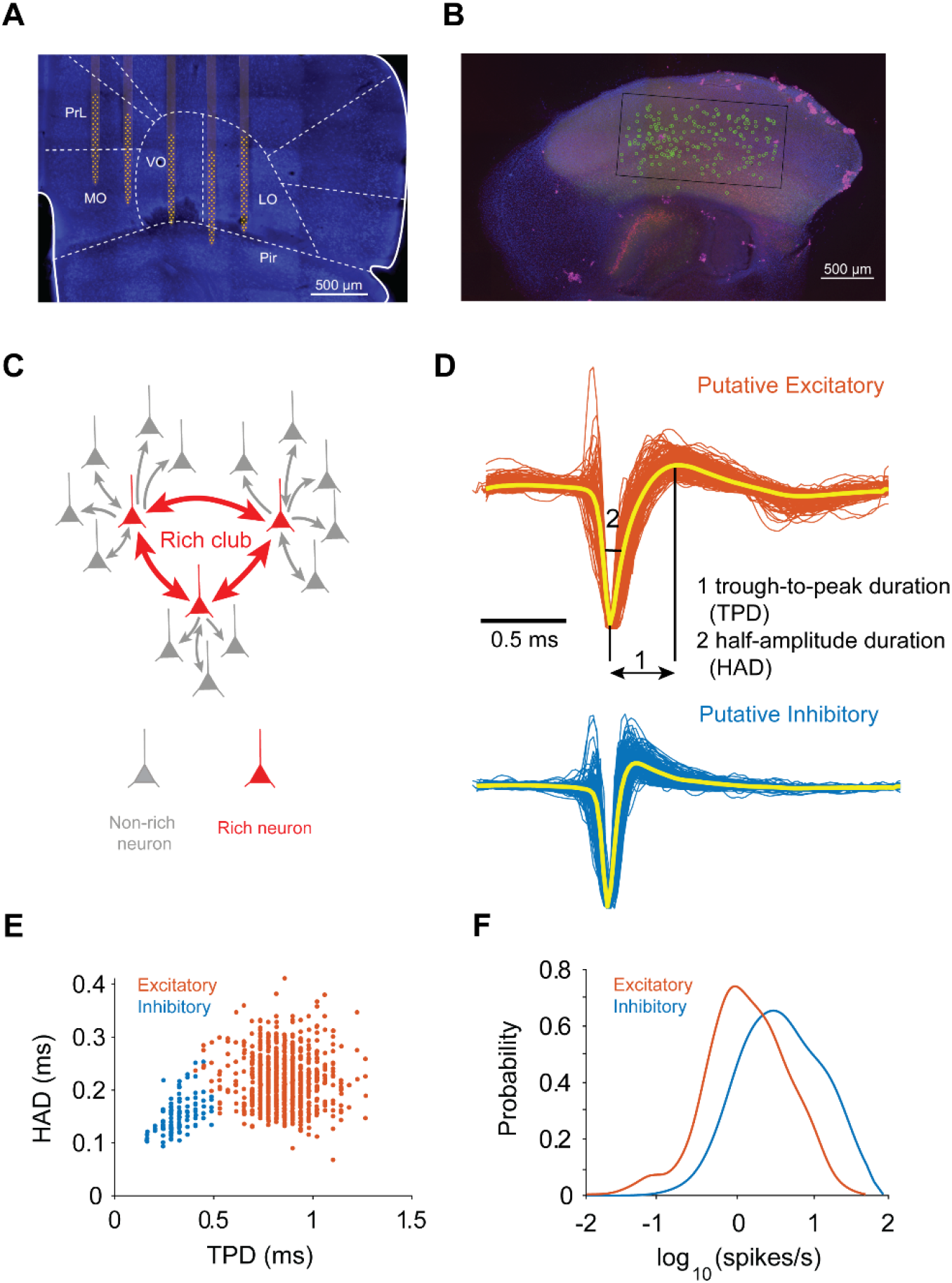
Experimental set-up, rich-club and cell type identification. ***A***, Spatial position of 5 silicon microprobes (256 electrodes in total) in orbitofrontal cortex and surrounding regions in awake head-fixed mice. Image shows a brain section with neuronal nuclei fluorescently labeled with a NeuN antibody (blue). The electrode positions were determined from tracks made by the silicon prongs visible in the sectioned tissue. For clarity, the five silicon prongs of the electrode array are superimposed on the image together with the recording sites (yellow). (PrL, Prelimbic cortex; MO, medial orbitofrontal cortex; VO, ventral orbitofrontal cortex; LO, lateral orbitofrontal cortex; Pir, piriform cortex). ***B***, Organotypic slice cultures placed on top of a 512 micro-electrode array (white rectangle). Yellow circles represent neurons identified after spike sorting. ***C***, Schematic of rich-club organization in neuronal networks, illustrating a few hub neurons (red) forming a dense and strongly connected subnetwork within themselves (red) and surrounded by a larger community of neurons (non-rich sub network) with fewer and weaker connections (gray). Thickness of the lines represents connection strength. ***D***, Spike waveforms of putative excitatory (orange) and inhibitory (blue) neurons from a representative session with trough-to-peak and half-amplitude duration marked; note the significant difference of spike width between the two putative cell types. ***E***, Scatter plot of spike waveform features from *D* showing two distinct clusters of neurons. ***F***, Firing rate distributions of putative inhibitory and excitatory neurons from the same session as in *D* and *E*.

### High concentration of inhibitory neurons within the rich-club

Do these functionally defined sub-networks differ in their cell-type composition? To examine this, we first used spike waveform features (trough-to-peak duration and half amplitude duration) and firing rates to classify single units as putative excitatory or inhibitory neurons (Ren et al., 2020) (see Materials and Methods) (Figure 1D). Cell types formed clearly separable clusters in feature space (Figure 1E) with distinct firing rate distributions (Figure 1F). Next, we isolated sub-networks for a range of normalized richness parameter values (*r*: sum of incoming and outgoing TE for each neuron, Figure 2A) and calculated the percentage of inhibitory neurons within these sub-networks. We observed a steady increase in the percentage of inhibitory neurons within sub-networks characterized by increasing richness parameter values (Figure 2B, D, Pearson correlation, r = 0.68, P < 0.001). We calculated chance levels of the percentage of inhibitory neurons in size matched sub-networks with the same richness parameters, by randomly permuting the identity of cell types throughout the network. Contrary to the actual networks with preserved cell identities, the percentage of inhibitory neurons in these permuted networks remained roughly uniform across the range of the richness parameter. Throughout the rest of the paper, the sub-network with normalized richness parameter of 0.8 (dashed vertical line, Figure 2B, D) will be defined as the rich-club (RC), although our findings were found to be qualitatively similar if we changed this threshold by ± 10%. Rich-club sub-networks were characterized by a higher percentage of inhibitory neurons compared to what was observed in the whole network (WN) both in-vivo (Figure 2C; Δinh_in-vivo_ = 12 ± 5%) and in-vitro (Figure 2E; Δinh_in-vitro_ = 23 ± 4%, Wilcoxon signed rank test, P < 0.001). In contrast, non-rich sub-networks had a lower percentage of inhibitory neurons compared to what was observed in the whole network (Δinh_in-vivo_ = -4 ± 1%, Δinh_in-vitro_ = -6 ± 1%; Wilcoxon signed rank test, P < 0.05). Our findings reveal the existence of inhibition-rich, densely connected sub-networks in both in-vivo and in-vitro cortical networks.

**Figure 2.**
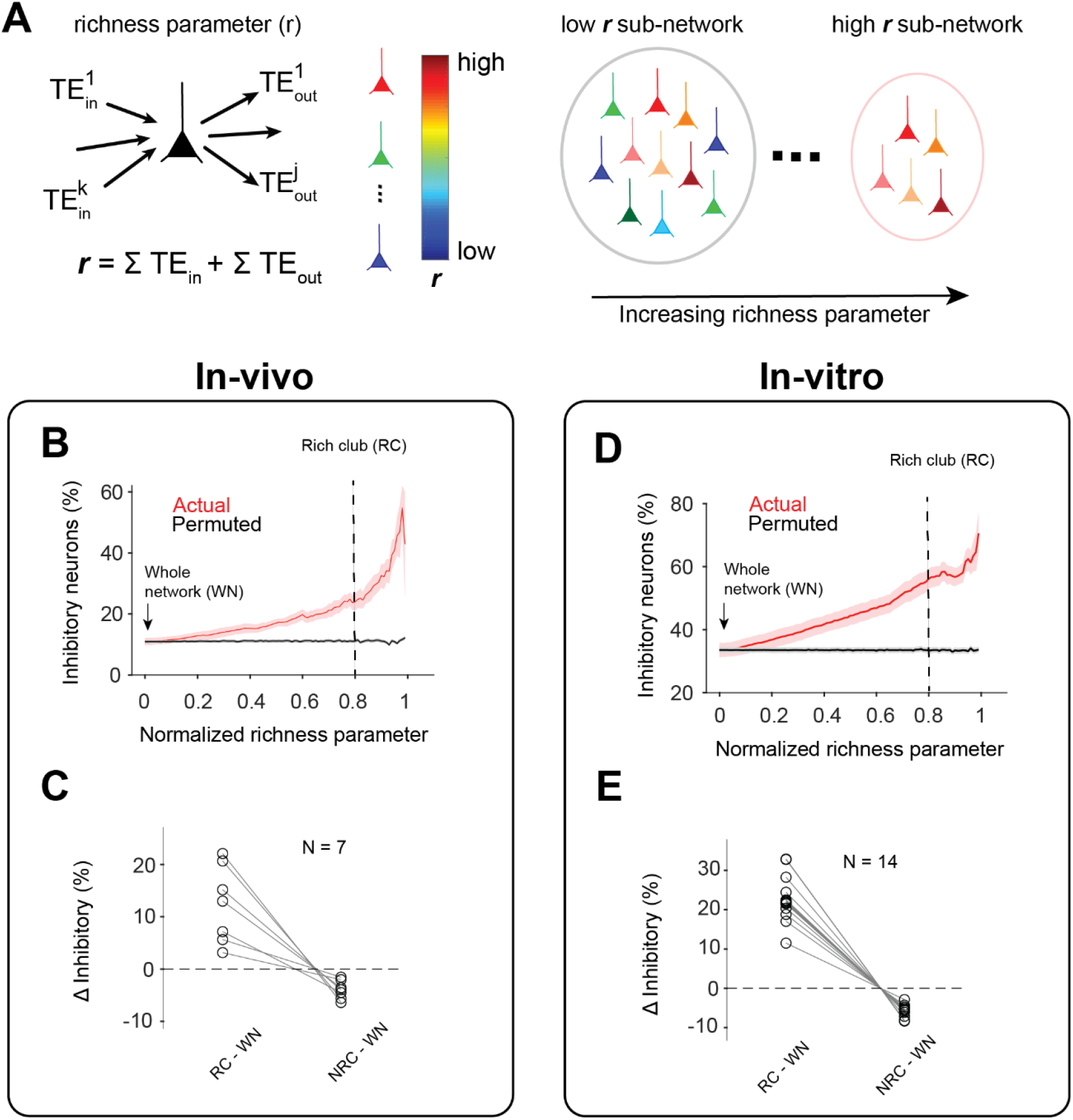
High concentration of inhibitory neurons in the rich-club. ***A***, Schematic showing how richness parameter (*r*) is defined for each neuron (left) and how different sub-networks are generated by setting increasing thresholds on the richness parameter (right). If the threshold is set at *r*_*1*_, then all neurons within the rich sub-network have richness > *r*_*1*_. ***B***, Percantage of putative inhibitory neurons (averaged across N = 7 in-vivo sessions) in subnetworks as a function of the normalized richness parameter (red) and for size matched networks with randomly permuted labels for excitatory and inhibitory neurons (gray; N = 100 permutations per data set). Solid curves represent the mean and shaded areas represent s.e.m. over all data sets. Vertical dashed line represents the threshold used to define the rich-club. ***C*, Left:** Difference in percentage of inhibitory neurons within the rich-club (RC) and the whole network (WN), **Right:** between the non-rich club (NRC) and whole network for N = 7 in-vivo sessions. Both positive and negative ΔInh values were significantly different from 0 (Wilcoxon signed rank test; P < 0.05). ***D-E*** same as in *B* and *C* except for in-vitro (N = 14) sessions.

### A highly synchronous group of inhibitory neurons within the rich-club

Given that inhibitory neurons are often implicated in synchrony (Bush and Sejnowski, 1996; Hasenstaub et al., 2005; Van Vreeswijk et al., 1994) in addition to synchrony’s established role in signal propagation (Bosman et al., 2012; Womelsdorf et al., 2007), we next sought to investigate whether the spiking dynamics within the rich sub-network showed more synchrony than the rest of the network. To quantify synchrony, we calculated the shuffle corrected Normalized Cross-Correlation Histograms (NCCH) between pairs of neurons within and across these sub-networks (see Materials and Methods). We analyzed a total of 73,228 pairs in-vivo and 712,366 pairs in-vitro across all sessions. Figure 3A shows NCCHs evaluated at different lags (-120 to +120 ms) for 3 example pairs with different sub-network memberships (inset shows zoomed-in plot with lags of ± 30 ms). To quantify pairwise synchrony, we evaluated the area under the NCCH curve between ± 30 ms and defined it as the Synchrony Index (SI; see Materials and Methods). The cumulative distribution of SI revealed network specific differences in synchrony (Figure 3B, cumulative distribution for an example session). Both in-vivo and in-vitro we found a higher percentage of pairs with SI values greater than 10% of the maximum SI (Figure 3C, F) indicating higher synchrony within the rich-club compared to the non-rich sub-network. To further examine the role of cell-types in synchrony in rich and non-rich sub-networks, we partitioned pairwise SI values into 3 classes: within inhibitory neurons only, within excitatory neurons only and across cell types (inhibitory and excitatory). We discovered that within the rich-club, inhibitory pairs had higher SI values compared to excitatory pairs or pairs with mixed cell types (Figure 3D, G). In contrast, within the non-rich sub-network only a small percentage of pairs exceeded SI values that were 10% of the maximum value observed in the networks (Figure 3E, H). Our findings indicate that network topology and cell-type jointly shape synchrony in cortical microcircuits.

**Figure 3.**
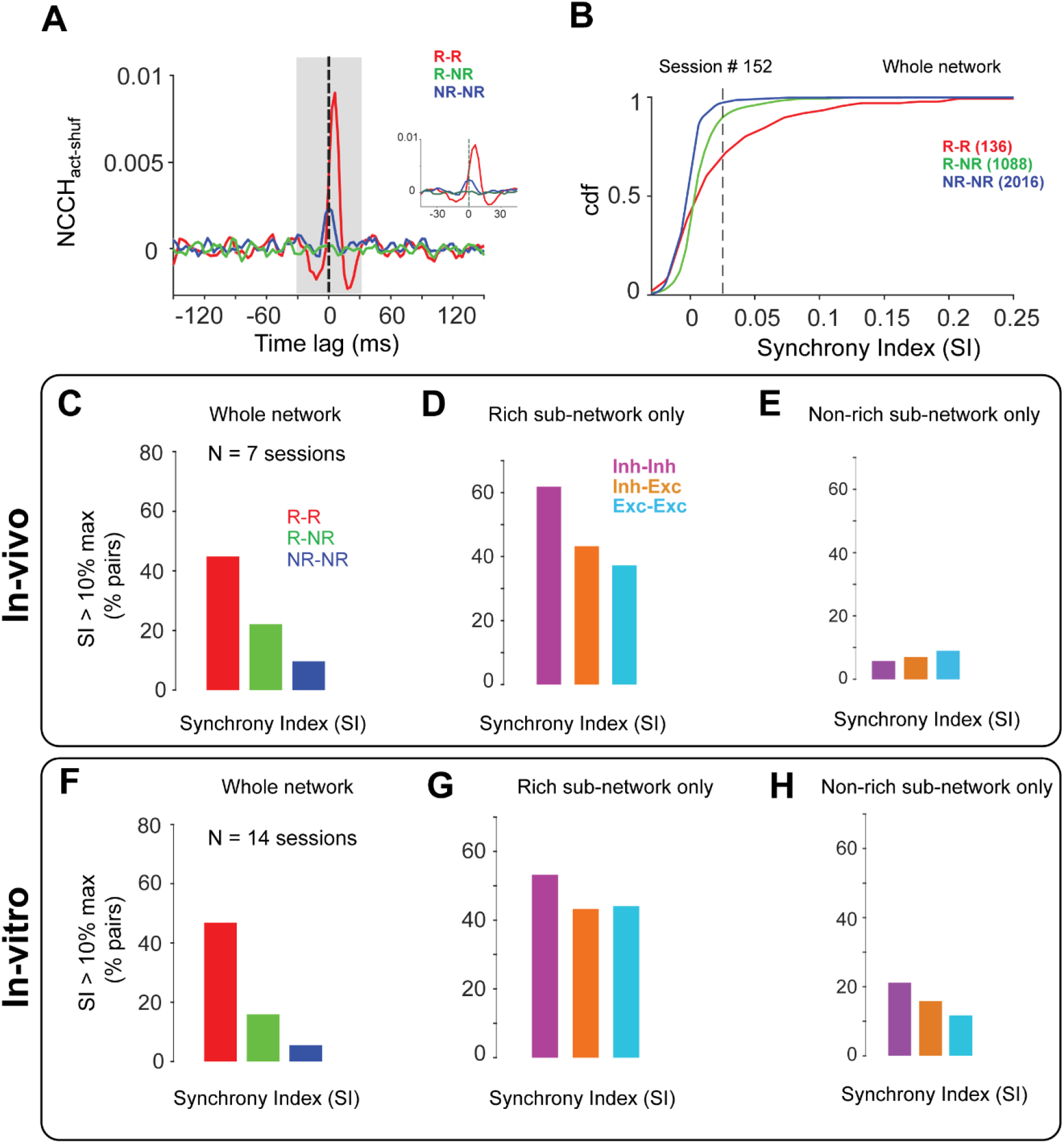
Higher pairwise synchrony of inhibitory neurons within compared to outside the rich-club. ***A***, Representative examples of shuffle corrected normalized cross-correlation histograms (NCCH) evaluated over a range of time lags (+120 to -120 ms), for pairs containing neurons only within the rich-club subnetwork (R-R, red), only within the non-rich subnetwork (NR-NR, blue) and a pair with mixed membership (R-NR, green). Gray shaded region (-30 to 30 ms) represents time-window used for the calculation of Synchrony Index (SI; see Materials and Methods). Inset shows a zoomed in version of the same NCCH between ± 45 ms. Note higher synchronous activity for the pair involving rich-club neurons compared to other pairs. ***B***, Cumulative distribution of SI for all pairs in an example in-vivo session, within and between rich and non-rich sub-networks as defined in *A*. Dotted black line represents 10% of the maximum SI value found in this session. Numbers in parentheses represent the number of pairs belonging to each class for the example sesssion. ***C***, Percentage of pairs for each class across all in-vivo sessions (N = 7) with SI greater than 10% of the maximum value. Note a higher percentage of inhibitory pairs compared to other pairs (roughly 2 times). ***D***, Same as in *C* except for pairs within the rich-club sub-network for different cell-type combinations (Inh-Inh, purple; Inh-Exc, orange; Exc-Exc, blue). E, Same as in *D* but for the non-rich sub-network. Note a larger percentage of inhibitory pairs with higher SI is only observed within the rich-club. ***F-H***, Same as in *C*-E except for in-vitro sessions (N = 14 sessions). Note cell-type specific effects on synchrony hold in both in-vivo and in-vitro networks.

### Rich-club shapes network dynamics

How do neurons within these distinct sub-networks influence activity at the population level? Single neurons within either sub-network (rich or non-rich) can trigger activity in the other through functional connections (Figure 4A). To examine this, we calculated the Population Normalized Cross Correlation Histogram (PNCCH) between the spiking activity of a single rich/non-rich neuron and the combined spiking activity of all the neurons in the non-rich/rich sub-network respectively at time lags extending from -150 to +150 ms (see Materials and Methods). PNCCH is an extension of the NCCH measure used earlier and captures more global associations (one to many) in spiking activity of neurons compared to NCCH (Okun et al., 2015). Figure 4B shows representative examples of PNCCH evaluated for the spiking activity of three rich neurons and the non-rich subnetwork. Heatmaps of PNCCH for all rich neurons from an example session show higher correlated activity post-spiking of a rich neuron compared to pre-spiking (Figure 4C). To quantify this further, we calculated the pre-spiking area under the PNCCH curve between -30 to 0 ms and the post-spiking area between 0 to 30 ms (Figure 4B, shaded regions). A population wide analysis showed that post PNCCH values were significantly higher compared to pre PNCCH values in case of both the rich (Figure 4D, G) as well as the non-rich neurons (Figure 4E, H) (Wilcoxon signed-rank test; P < 0.01). However, the difference between post and pre-values (ΔPNCCH) was significantly higher for the rich neurons compared to the non-rich neurons (Figure 4F, I) (Wilcoxon rank-sum test; P < 0.001). Our results indicate that there is an asymmetry in how single cells impact network activity and this is likely due to the difference in locations of the neurons in the overall functional network topology.

**Figure 4.**
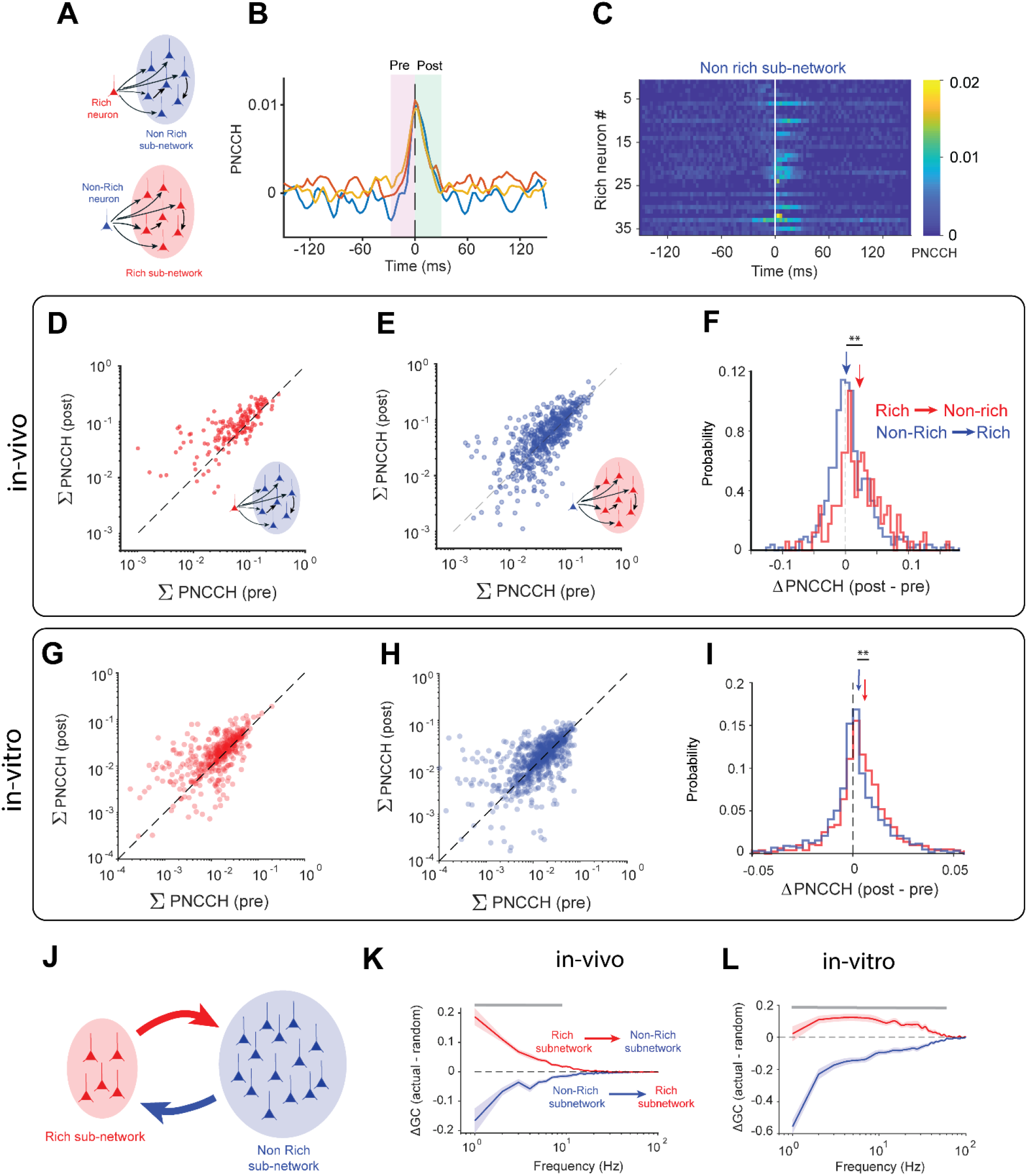
Rich club neurons shape network activity in cortical networks. ***A***, Schematic representation of how single neurons within the rich/non-rich subnetworks can influence activity in other subnetworks through functional interactions. ***B***, Population normalized cross-correlation histogram (PNCCH) for three rich neurons evaluated for a range of time lags. Shaded regions represent time windows pre- and post-spiking of the reference neuron used for further analysis. ***C***, Representative PNCCH for all rich neurons from an example session. Horizontal rows in the heat map represent the PNCCH of the spiking activity of a single rich neuron and the population activity of non-rich neurons calculated at different time lags (-150 to +150ms) (see Materials and Methods). Note higher values of PNCCH post-spiking of a rich neuron compared to pre-spiking. ***D***, Summed post (0 to +30 ms) and pre PNCCH (-30 to 0 ms) values for all rich neurons across all 7 in-vivo sessions (red filled circles; N = 188 neurons). Inset shows the reference population (non-rich sub-network) used to calculate the PNCCH. Same as in *D* except PNCCHs are evaluated between the spiking activity of each non-rich neuron (blue dots; N = 435 neurons) and the combined spiking activity of neurons in the rich-club sub-network (inset). ***F***, Distribution of the difference in summed PNCCH values pre- and post-spiking for rich neurons (red) and non-rich neurons (blue). Blue and red arrows represent the median values of each distribution (asterisks denote statistical significance of a Wilcoxon rank-sum test; P<0.001). ***G-I***, Same as in D-F except for in-vitro sessions (N = 14 sessions). The number of rich and non-rich neurons analyzed to calculate the PNCCH values for the in-vitro sessions were 916 and 2066 respectively. ***J***, Schematic showing how overall population activity in each subnetwork can drive activity in the other through functional interactions. ***K***, Difference of Granger causality values between actual and size matched randomly sampled networks (see Materials and Methods) as a function of frequency from rich to non-rich sub-network (red) and vice versa (blue). Solid lines represent mean values across N = 7 sessions and shaded regions represent s.e.m. Gray bar represents frequencies for which ΔGC is significantly different from 0 (signed rank test followed by Bonferroni Holm’s multiple comparison test) ***L***, Same as in *K* except for the in-vitro sessions.

To further analyze the strength and directionality of the interactions between the rich and non-rich sub-networks at the scale of neuronal populations, we estimated Granger causality (GC, see Materials and Methods) from rich to non-rich and non-rich to rich sub-networks using the population spiking activity in these sub-populations (Figure 4J). To specifically examine the role of network topology, we randomly sampled neurons from the entire network to create size-matched pseudo rich and non-rich sub-networks and calculated GC values between these pseudo sub-networks. We then subtracted these values from the estimate of GC between the actual rich/non-rich sub-networks (ΔGC). The rich club sub-network exerted higher than expected Granger causal influence on the rest of the network (ΔGC>0; signed rank test, Bonferroni-Holm multiple comparisons test) whereas the Granger causal influence of the non-rich club-network was significantly lower than expected by chance (ΔGC>0; signed rank test, Bonferroni-Holm multiple comparisons test; Figure 4K, L). Our results suggest that network topology endows the rich-club with a greater capability to influence network activity in other parts of the network.

### Network topology shapes avalanche dynamics

Finally, we examined to what extent network topology influenced signal propagation in these networks. We investigated neuronal avalanches as indicators of signal propagation. Previous work with multielectrode array recordings (Beggs and Plenz, 2003) defined a neuronal avalanche as continuous spiking activity spanning one or more time bins, bracketed by time bins of no activity at the beginning and at the end (Figure 5A). The number of consecutive time bins with at least one active neuron is defined as the avalanche length. We first calculated the mean pairwise synchrony (SI) between all pairs of neurons involved in short (L < 15 ms) and long avalanches (L > 15ms). Longer avalanches exhibited significantly longer tailed distributions of SI compared to shorter avalanches (Figure 5B, C; 2-sample KS test, P < 0.05) indicating that highly synchronous pairs were more likely to be present in longer avalanches compared to shorter ones. Since we previously showed that the rich-club had higher synchronous activity compared to the non-rich sub-network, we examined how the participation of the rich neurons at each time step of an avalanche scaled with the length of the avalanche (example session, Figure 5D). We found highly significant positive correlations (Pearson’s correlation) between the avalanche length and the mean fraction of rich neurons participating in those avalanches (Figure 5D inset). Across all sessions we observed this trend in 5 out of 7 in-vivo sessions and 10 out of 14 in-vitro sessions (Figure 5E). Our findings indicate that a highly synchronous sub-network of rich neurons influences the spread of spatiotemporal patterns in cortical microcircuits both in-vivo and in-vitro.

**Figure 5.**
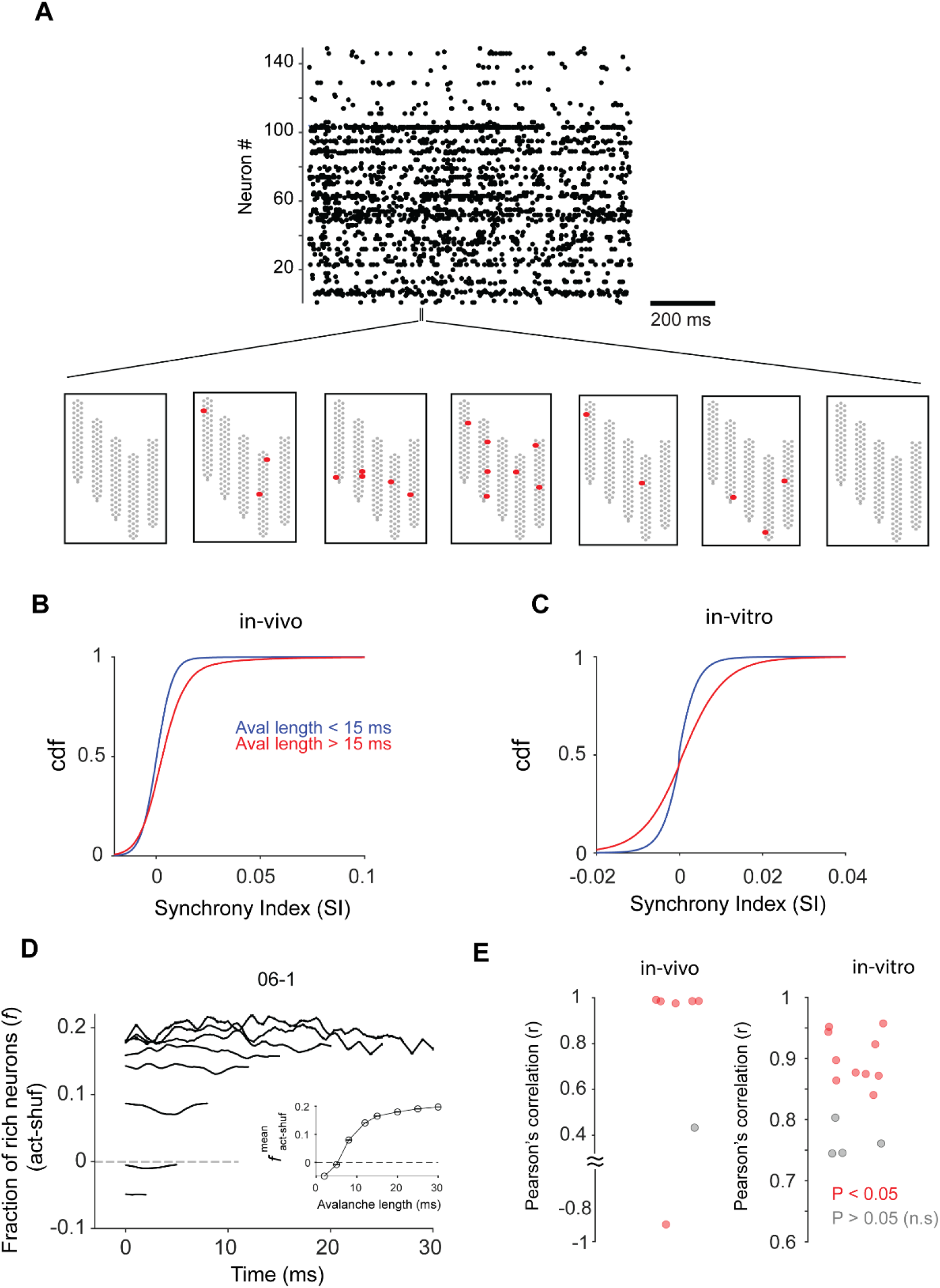
Fraction of rich-club neurons active in avalanches increases with length. ***A***, Example spike raster of simultaneously recorded neurons (N = 142) from an example session (top). Representative example of a neuronal avalanche: a spatiotemporal sequence of neuronal activity with at least one active neuron per time bin, flanked by time‐bins with no activity (bottom). Gray dots represent electrode locations on the silicon probe and red dots represent active single units at each time step of the avalanche. ***B***, Cumulative distribution of SI for all pairwise combinations of neurons participating in short (< 15 ms; blue line) and long avalanches (>15 ms; red line). ***C***, Same as in *B* but for in-vitro data. ***D***, Shuffle corrected fraction of rich club neurons (see Material and Methods) active in different time steps of avalanches of various lengths for an example session. Inset shows the mean value (averaged across all time points) of the fraction of active rich neurons in avalanches of various lengths. ***E***, Pearson’s correlation between mean fraction of active rich neurons and avalanche length for individual sessions in-vivo (left) and in-vitro data (right). Each filled circle represents a session and the color represents the p-value of the correlation (red:P < 0.05, gray: P > 0.05).

## Discussion

The dynamics in networks of cortical neurons is expected to be strongly influenced by cell type composition and the pattern of connections between them (Bojanek et al., 2020; Jabri and MacLean, 2022). Here we show that within subnetworks (in-vivo and in-vitro) consisting of highly connected hub neurons (rich-club), the abundance of inhibitory neurons is significantly higher compared to that in the entire network or the non-rich subnetwork. Inhibitory neurons within the rich-club exhibit higher pairwise synchrony compared to other pairs within and outside the rich-club. The rich club showed stronger than expected Granger causal influence on the rest of the network at a broad range of frequencies. Finally, the participation of rich club neurons in avalanches increased with the length of such spatiotemporal patterns suggesting that they could play an important role in the propagation of network wide activity. Our findings provide a novel perspective on how cell type composition (excitatory vs inhibitory) coupled with a non-random network topology (highly connected rich-club) plays a central role in regulating network dynamics in cortical microcircuits. Interestingly, a similar finding emphasizing that cell type and network topology jointly determine network dynamics has recently been reported for *C. elegans* (Uzel et al., 2022).

### Relation to previous work

Previous studies in the hippocampus have shown the existence of densely connected neuronal hubs which were primarily GABAergic interneurons (Bonifazi et al., 2009; Picardo et al., 2011) and played a crucial role in regulating overall network synchronization (Bonifazi et al., 2009). However, whether densely connected hub neurons were present in cortical circuits and to what extent they influenced network activity was unknown. We address these questions by showing that in cortical circuits both in-vivo (orbitofrontal cortex of awake behaving mice) and in-vitro (organotypic cultures), sub-networks of highly connected hubs (rich-club) have a higher than expected abundance of inhibitory neurons. Although there are differences in how neural circuits evolved in the intact brain compared to cultures, our results indicate that there is an overarching general theme to the organization of neural circuits (cell-type specific connectivity), across brain areas and preparations. In fact, in a recent study Kajiwara and colleagues (Kajiwara et al., 2021) found that inhibitory neurons were more centrally located within the functional network topology of somato-motor cortex in mouse slices.

### Linking inhibition to activity propagation

We observed that although the rich club has a relatively higher concentration of inhibitory neurons, it exerts Granger causal influence on the rest of the network and shows higher activation levels for longer avalanches. It would seem paradoxical that the rich club containing a higher than expected concentration of inhibitory neurons is central to the propagation of longer spatiotemporal patterns. However, characterizations of connectivity patterns within different classes of inhibitory neurons in the visual cortex (Pfeffer et al., 2013) and electrical and chemical synapses between inhibitory neurons in the cerebellum (Rieubland et al., 2014 have shown that inhibitory neurons can inhibit each other. This could potentially lead to the disinhibition of excitatory neurons that are connected to such populations through overrepresented three neurons motifs, for example I→I→E (Gal et al., 2017; Shimono and Beggs, 2015; Song et al., 2005) along with other motifs such as I→E→I and E→I→I. Recent work on chemogenetic activation of inhibitory neurons has shown that such activation suppresses the activity of most interneurons in addition to the expected suppression of excitatory pyramidal cells (Rogers et al., 2021). Interestingly, a computational model of working memory predicts that enhancing I→I connections would lead to more stable dynamics (Kim and Sejnowski, 2021). This would be consistent with recordings from zebra finches showing that the presentation of songs over many days is accompanied by a stable pattern of inhibitory activation, while the population of excitatory neurons is less stable (Liberti et al., 2016). Future studies can perform more detailed characterizations of connectivity within and across excitatory and inhibitory populations to further tease apart their differential roles in network dynamics in neuronal populations.

### Limitations of current work

Although our findings show that inhibitory neurons within the rich club play a crucial role in regulating network dynamics, the specific cell morphology of these neurons is not known. Hub neurons in the hippocampus have been found to consist of two types: GABAergic interneurons with long-range axonal projections and basket-like neurons with dense local arborizations (Bonifazi et al., 2009). Interestingly, phasic stimulation of only the basketlike hub neurons led to network synchronization. It was hypothesized that basket cells could act as local hubs whereas long axon projecting hub interneurons could play the role of connector hubs. In the neocortex inhibitory neurons are broadly grouped into PV+, SST and VIP (Tremblay et al., 2016), although recent work has shown that in the visual cortex there are at least 15 different types of inhibitory neurons with distinct morphological and electrophysiological properties (Jiang et al., 2015). Characterizing the morphology of inhibitory hub neurons and examining specific classes of inhibitory neurons within the cortical rich-club will add further insight into how cell type differentially regulates local and long-range network dynamics.

Another potential limitation of this work is that the structural connectivity (synapses, gap junctions) of the neurons from which we record is unknown. Despite impressive technical advances in connectomics, to the best of our knowledge only a few studies have created a complete reconstruction of a local cortical network using electron microscopy (Turner et al., 2020; Yin et al., 2020); it is much more common to instead provide statistical descriptions (Erö et al., 2018). While it would be extremely desirable to obtain the connectome of such a local cortical circuit, this would not necessarily reveal its dynamics, in the same way that a road map by itself would not definitively indicate traffic flow. A common example in this regard is the case of *C. elegans*, where the connectome has been known for decades (White et al., 1986), yet fruitful work to model how this network routes activity and influences behavior is still revealing surprising new findings (Izquierdo and Lockery, 2010; Luo et al., 2014; Randi and Leifer, 2020). Functional connectivity, like the TE networks we construct here, has proven to be extremely useful at the whole brain level in distinguishing between health and disease (Lynall et al., 2010, Hadjiabadi et al 2021), conscious and unconscious states (Achard et al., 2012), and networks that have learned from those that have not (Bassett et al., 2011). Given this promising record, we expect that studying functional networks of cortical neurons, like we do here, will be an important step toward identifying microcircuit changes that underlie disease states and learning (Schröter et al., 2017).

### Rich club networks and sensory coding

What could be the possible role of this distinct subnetwork in encoding sensory information or behavior in cortical networks? Specifically, do these subnetworks have a greater capacity of encoding or decoding sensory/non-sensory information? Previous work in cortical columns in macaque V1 has shown the presence of synergy hubs i.e., neurons that engage in predominantly synergistic interactions with other neurons to encode stimulus information (Nigam et al., 2019). Interestingly, sub-populations consisting of only synergy hubs were better at decoding stimulus information compared to redundancy hubs. However, the cell type composition of synergy hubs and whether they form a densely connected rich-club is unknown. Additionally, recent work in cortical slice cultures has shown that neurons with the highest synergy values tend to reside in the rich-club (Faber et al., 2019). Future work in sensory as well as non-sensory cortical areas can further explore these missing links between network topology, cell type composition and information encoding in neuronal populations.

### Implications for cortical models

Our findings have important implications for cortical network models. Traditionally, these models have used an 80/20 rule for determining the size of excitatory and inhibitory populations. The finding that distinct subnetworks deviate from this excitatory to inhibitory ratio poses the question whether a more compartmental model with different abundances of inhibitory and excitatory neurons and connectivity profiles needs to be implemented. Such models can also provide a more detailed understanding of the source and propagation of activity patterns by selectively perturbing different cell types within such networks (Sadeh and Clopath, 2020). Conversely, cortical models could explore what plasticity mechanisms could lead to the formation of an inhibition-dominated, strongly connected rich-club starting from a randomly connected network topology.

## Statistical analysis

We have used non-parametric statistical tests throughout the manuscript as mentioned in the main text. Specifically, for comparing cumulative distributions with different sample sizes we have used the 2-sample KS test. In cases where comparisons were made between 2 distributions with pairwise correspondence, we used the Wilcoxon signed-rank test.

## Acknowledgements

This work was supported by a Robust Intelligence grant (NSF # 1513779) and IU OVPR bridge funding from Indiana University to JMB, Whitehall Award (ID:1712-114) and IU OVPR Bridge funding from Indiana University to ELN, NIH R01 (grant # 1R01MH121978 to OS and NSF grant (# 1651396) to IHS.

This research was supported in part by Lilly Endowment, inc., through its support for the Indiana University Pervasive Technology Institute. The authors also acknowledge the Indiana University Pervasive Technology Institute for providing supercomputing and storage resources.

## Data and code availability

Custom written software used for the analysis reported in this study is available at https://github.com/hadihafizi/InhibRichClubDyn. In-vivo spike data are available at https://github.com/sotmasman/Cortical-dynamics and in-vitro data can be accessed from the Collaborative Research in Computational Neuroscience (CRCNS) data sharing website at http://dx.doi.org/10.6080/K07D2S2F.

## Author contributions

HH and SN performed the analysis and wrote the manuscript. JB performed granger causality analysis and edited the manuscript. NR and HIS performed cell type classification and edited the manuscript. SCM, OS and ELN edited the manuscript. JMB wrote the manuscript and conceived and supervised the project.

